# Motor adaptation to environment changes predicting interactable object behaviour can be flexible and implicit

**DOI:** 10.1101/2024.02.06.579172

**Authors:** Shanaathanan Modchalingam, Andrew King, Bernard Marius ’t Hart, Denise Y. P. Henriques

## Abstract

The human motor system can adapt to perturbations by updating existing models of motor control or by creating context-specific motor memories or strategies. We investigate if motor adaptation is context-informed when a perturbation is applied to either the throw direction of a ball, or the acceleration of the ball post-release during a virtual throw-to-target task. Using the visual slant of the task surface in an immersive virtual environment, we determine if the tendency for model updating is influenced by informative visual cues that predict perturbations to intended actions. Our findings reveal that perturbations resembling accelerations enabled flexible motor adaptation regardless of the presence of the slant cue. Perturbations in the throw direction conversely led to internal model updating. Additionally, visual slant properties of the task surface elicited implicit, slant-specific changes in performance. Our findings underscore the role of visual properties of both perturbations and environments in flexible motor learning.

## Introduction

The ability to flexibly adapt to changing environments is crucial for successful performance in many real-world motor tasks. Upper limb motor adaptation has been extensively studied in the past by perturbing the hand movement or representations of the hand, leading to errors in predictions of sensory consequences of movements and eliciting an updating of internal models governing actions. Although such motor adaptation paradigms well represent internal changes to the body, such as injury or fatigue, a significant portion of motor learning in daily activities involves the motor system adapting to changes in objects and the surrounding environments. When adapting to these externally localized perturbations, there may be clear visual indicators that predict the behaviour of objects of interaction, such as the movement of grass on a golf course that predicts the strength of a sidewind. In such situations, rather than updating current model for the task, motor adaption may rely on forming context-specific motor memories or cognitive strategies, allowing for effective and rapid task switching when environments change.

Understanding how the brain adapts to real-world perturbations is essential for developing effective interventions and training programs for individuals with motor impairments or those seeking to improve their motor skills. In this study, we tested if adaptation to perturbations that affect the acceleration of a thrown object, suggesting external sources of errors, leads to more flexible motor adaptation than the internal-model updating expected when adapting to visuomotor rotations, which suggest internal sources of error. Although participants adapted to both types of perturbations, those adapting to accelerations in ball paths and even perturbations visually resembling accelerations did not show evidence for model updating. Instead, these perturbations facilitated more flexible motor learning. We additionally tested if immersive visual cues that could plausibly explain errors or aid in strategy-formation could supress model updating. We find informative changes in the visual environment did not affect model updating characteristics but could cue immediate and implicit changes in task performance during learning and de-adaptation.

In prior work, systematic differences between predicted and actual consequences of planned upper limb movements reliably lead to the updating of internal models of motor control that map goals to actions [1–4]. Updating existing models avoids switching costs associated with motor memory selection and retrieval, allowing for efficient and timely movements [5,6]. Motor errors can be reduced exponentially via one or more learning processes [7–10]. However, this approach is not ideal when motor contexts change rapidly as these models must be re-updated to previously experienced contexts through the same learning processes, and errors must again be decreased exponentially over time. More flexible motor learning, either through the creation of new implicit motor memories or through strategy formation, may improve task-success when movement types, objects of interaction or environments change rapidly [11]. Although behaviour patterns indicating model updating have been observed when adapting to errors predicting a movement or hand-object interactions [12–15], the relative contributions of model-updating or more flexible motor learning when adapting to errors in predicting object-environment remains unclear. In this study, we use a target-directed ball throwing task to test if motor adaptation patterns differ when adapting to perturbations that are likely due to environment interactions, like accelerations added to the movement paths of thrown objects, or hand-object interactions, like rotations of intended throwing directions.

Localizing the source of a given error during a motor task can be crucial for determining if existing internal models should be updated or if an alternative model or strategy should be created. Perceptual cues the suggest a change in movement type, like changes in hand or body posture, reliably lead to flexible motor adaptation through the formation of new motor memories that can be switched to when returning to previously experienced conditions [16–19]. When the movement type remains consistent, findings on the efficacy of interactable object or environment-based cues to facilitate flexible motor adaptation are mixed [20–25]. Generally, properties extrinsic to movements, such as the colour of objects in the environment or object identity do not facilitate the creation of new motor memories [18,20,23]. In some cases however, visual properties of the environment may serve to explain errors in outcomes under a given predictive model and render model updating less useful [26], allowing for flexible motor learning, possibly through the use of explicit strategies [12,15,24,25,27,28]. Both saliency and task relevance are crucial for detecting changes in context and may explain these conflicting findings [29,30]. Although cues like environment colour are salient, they may not facilitate context-change detection to the same degree as task-relevant cues [12,29,31]. In dual-adaptation paradigms for example, although the colour of the target of reaching movements may fully indicate the presence of a target-colour-specific perturbation, the creation of target-colour-specific motor memories is not likely since the colour of external objects is usually irrelevant to internal models of upper limb movements. It is not clear however, if this holds when visible environmental changes fully predict changes in object-environment interactions. In studies where motor learning intrinsically involves learning the interaction between movement and the environment, properties of the object being interacted with are essential, as learning is often optimized to account for object or environment properties even when errors are low [32,33]. Here, we examined whether adaptation patterns differ from those in upper-limb motor adaptation tasks when errors could be explained by object-environment-interactions in an immersive virtual environment. Specifically, we investigate if changes in internal model updating caused by our prescribed internally and externally attributed perturbation types is influenced by visual cues in the environment that reliably predict the behaviour of thrown objects, such as the slope of a surface on which thrown objects travel.

In a series of experiments, participants made realistic throwing movements to hit targets in a virtual task where the thrown object rolled on a flat surface. During the experiments participants adapted to two perturbation types: visuomotor rotations at the moment of release, often internally attributed, and perturbations resembling real-world accelerations, acting on the object after the release. We test if acceleration-like perturbations lead to flexible motor adaptation allowing for fast changes in performance in new or repeated motor contexts. We further test if immersive and informative visual cues, which may prompt faster motor adaptation, can also lead to more flexible motor learning for both internally and externally attributed perturbation types. We find that accelerations, and even perturbations that are visually acceleration-like, can lead to more flexible motor learning than model updating. In all cases, immersive visual changes to the slant of the task surface prompted immediate behaviour change but did not facilitate the creation of new motor memories or strategies required for flexible motor learning. In a subsequent experiment, these visual environment-cued behaviour changes persisted even while participants supressed cognitive strategies during throwing movements. We conclude that behavior changes triggered by informative and task relevant changes in the environment can occur implicitly. This implies that the updated models in these instances include models of ball-environment interactions, rather than models specifically of the arm or release dynamics.

## Methods

### Participants

169 participants (103 female, 20.75 ± 4.55 years of age, mean ± SD) participated in the study and were divided into one of 8 groups within 4 experiments. All participants had normal or corrected-to-normal vision and were naïve to visuomotor learning experiments. All participants voluntarily participated in the study and provided written and informed consent. The procedures used in this study were approved by York University’s Human Participant Review Committee and all experiments were performed in accordance with institutional and international guidelines.

### Apparatus/Setup

Participants sat on a height-adjustable chair with a table at waist height. Armrests were adjusted to match the table height and participants rested their elbows on the armrests for comfort between trials. Participants were then verbally instructed on the details of the task. The instructions used can be found at this project’s Open Science Framework repository (osf.io/a5nv3/). After receiving the verbal instructions, participants donned a head-mounted display system (HMD: Oculus Rift Consumer Version 1; resolution 1080 by 1200 for each eye; refresh rate 90 Hz) and grasped Oculus Touch controllers in both hands. Three Oculus Constellation sensors tracked the positions of the HMD and Oculus Touch controllers. Participants received all visual feedback via the HMD. The virtual-reality environment was developed in Unity 3D, using the Unity Experiment Framework to handle trial and block schedules during the experiment [34]. Before starting the experiments, participants performed practice trials where they were encouraged to explore the tasks involved in the experiment. Participants could repeat the practice trials if required. Once all practice trials were completed, further instructions were provided by the experimenter and participants began the experiment.

### Roll-to-Target Task

In all experiments, participants repeatedly performed the Roll-to-Target task. The goal of the task was to roll a ball such that it travelled as close as possible to a target location (Fig 1a). Participants received a score from 0-10 for each roll based on the minimum distance between the ball path and the target. Participants also received +5 bonus points for accurately hitting the target.

**Figure 1:**
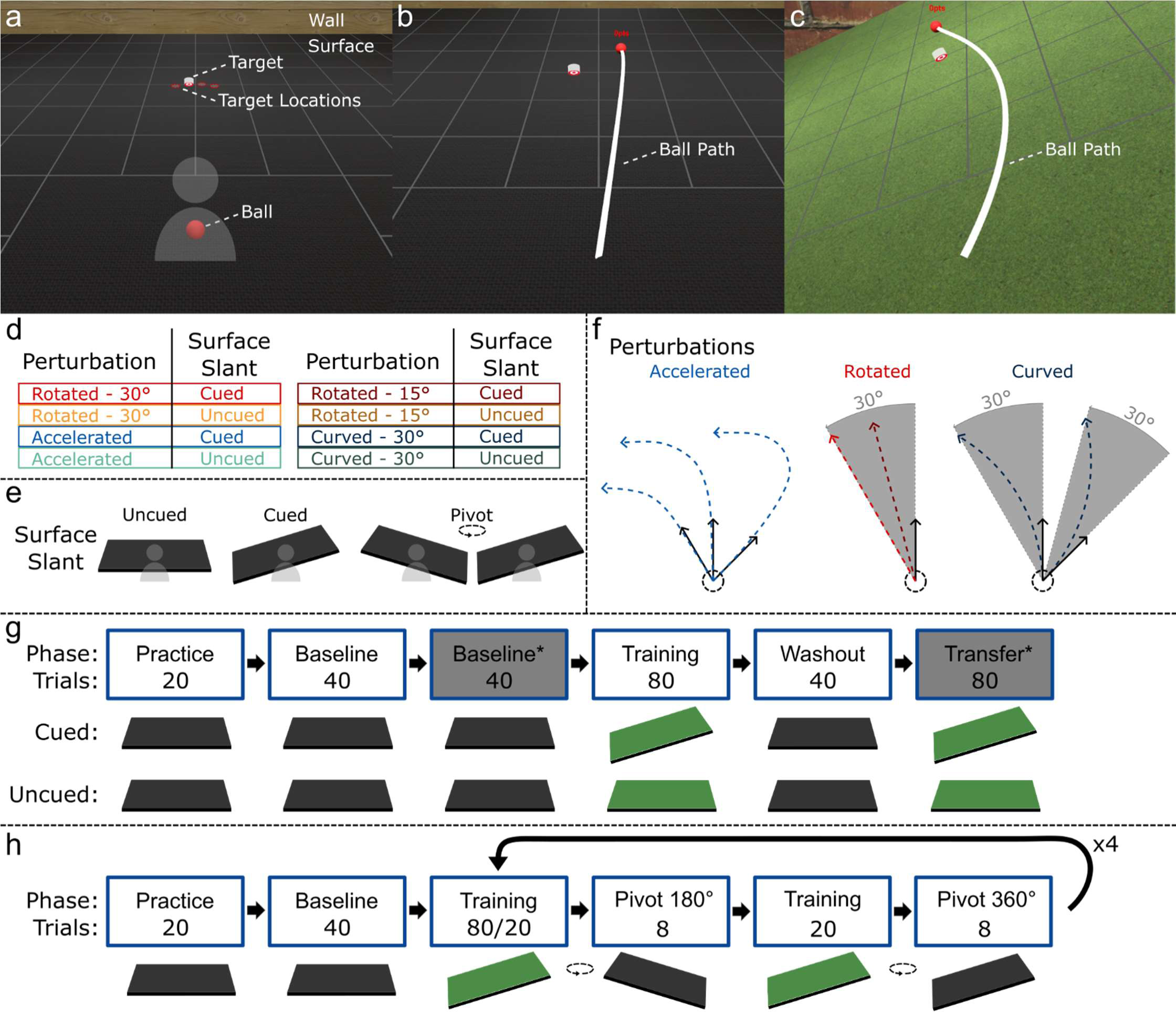
**a**: Visual display in HMD at the start of each trial in the Roll-to-Target task. The target appeared at one of 4 possible target locations. **b**: Visual display at the end of the Feedback phase of a Roll-To-Target task with the task surface aligned with the real-world floor and Baseline-accelerations. **c**: Visual display at the end of the Feedback phase of a Cued-Curved Roll-to-Target task. **d**: Possible perturbation/cue combinations during the Training and Transfer phases of Experiment 1. **e**: Possible immersive visual slants of the task surface. During Cued Roll-to-Target tasks in Experiment 1, the task surface was slanted 25°. In Experiment 2, a 25° visual slant acted as the cue during the Training phase and the animation prior to each Washout phase resulted in a -25° or 25° visual slant for 180° and 360° animations respectively. **f**: Possible perturbations during the Feedback step of Roll-To-Target tasks. **g**: Phases and trials in Experiment 1. Surface Baseline* and Transfer* phases were not analyzed in this study. Participants used their non-dominant hands during Baseline* and Transfer* phases. **h**: Phases and trials in Experiment 2.

Each trial of the task began by participants holding their right hand near the midline of their body below the chin. In the virtual environment, they were situated over a large horizontal plane (4 m x 4 m) located at waist height, termed the Surface (Fig 1a). A wall with a horizontal wood plank or brick texture at the far end of the Surface, along with grid-lines along the task surface, provided an accurate visual representation of the orientation of the Surface. A virtual 10-cm diameter ball was placed in front of the participant at the “starting position”. A target was placed 1 meter away from the starting position of the ball in one of 4 possible locations on the Surface (84°, 88°, 92° and 96° in polar coordinates: Fig 1a). Target locations were pseudorandomized so that in every 4-trial set, participants threw to each target at least once.

The goal of the Roll-to-Target task was to make a throwing movement to roll a ball and hit targets as accurately as possible. Each trial contained an Action step and a Feedback step. Each trial began with an Action step, where participants made a throwing movement to the target by holding down the trigger button on the Oculus Touch controller with their index finger and making a quick, outward motion with their dominant hand. The ball was released when the hand moved 15 cm from the starting position of the throw, or when the trigger button was released, whichever event occurred first. To ensure consistent rolling motions, participants needed to complete the Action step within 1.5 seconds from the start of the trial to receive a score for the trial. Although participants did not receive a score under this time-out condition, we included these trials in our analysis. The Feedback step began immediately upon the release of the ball.

During the Feedback step, participants observed the ball roll and were given feedback on the ball path. The speed and direction of the throwing movement at the point of release in the Action step determined the speed and travel path of the ball in the Feedback step. The speed of the ball was capped at a maximum of 3.96 m/s. The feedback step ended either after hitting the target or after 2 seconds had elapsed from the start of the Feedback step. At the end of the Feedback step participants were shown a visual representation of the ball path and received a score for the trial based on the minimum distance between the ball path and the target (Fig 1b-c).

### Task Variations

We used 4 variations of the Roll-To-Target task that modified the relationship between the throwing movement in the Action step and the ball path in the Feedback step: Aligned, Rotated, Accelerated, and Curved (Fig 1d). While all participants performed Aligned Roll-To-Target tasks, the Rotated, Accelerated, and Curved variations were considered perturbations and each participant only experienced one of the 3, dependent on their group (see Experiment Protocol).

In Aligned variations of the task, the visual Surface was aligned to the real-world floor-plane. In all other variations, the Surface was either in the same visual alignment as the Aligned variation, or displayed with a 25° CCW visual slant (Fig 1c) in groups termed “Uncued” and “Cued” respectively (Fig 1d-e).

In Aligned Roll-to-Target tasks, the direction and speed of the throw in the Action step determined the initial heading direction and speed of the ball in the Feedback step. In Aligned tasks, we continuously applied ‘Baseline Accelerations’ to the ball path (-0.31, -0.15, 0.15, 0.31 m/s^2^, pseudorandomized). Baseline Accelerations were used so that all Roll-to-Target trials involved changes in the ball’s direction of travel. Baseline accelerations were small, centered around zero and had an unpredictable order to have little to no effect on task performance.

In Accelerated Roll-to-Target tasks, the initial direction of the ball movement during the Feedback step matched the throw direction in the Action step, while a constant leftward acceleration of 2.49 m/s^2^ was applied to the ball as it rolled during the Feedback step (Fig 1f). The movement path of the ball during this task variation resembled a plausible movement path on a surface with a 25° surface slant. Although all Roll-to-Target tasks included accelerations in the ball path during the Feedback step, only the acceleration in Accelerated tasks could be reliably countered.

In Rotated Roll-to-Target tasks, the speed of the throw in the Action step determined the initial speed of the ball, but the initial heading direction of the ball in the Feedback step was rotated by 30° or 15° counter-clockwise relative to the throw direction along the Surface plane (Fig 1f). We applied Baseline Accelerations to all Rotated Roll-to-Target tasks. We used a rotation size of 30° and an acceleration of 2.49 m/s^2^ in the experimental groups to match the changes in throw angles required to reliably hit targets with Rotated and Accelerated perturbations. In a follow up experiment, we used a rotation size of 15° to match the error sizes participants observed in the Accelerated perturbations.

In Curved Roll-to-Target tasks, the initial speed and direction of the ball movement during the Feedback step matched the throw speed and direction during the Action step. During this task variation, the path of the ball curled in a counter-clockwise direction so that the ball passed through a point 30° counter-clockwise to the throw direction when at the target distance (Fig 1f). The magnitude of the angular offset of the ball path was scaled to the distance from the starting position of the ball. Curved perturbations required the participants to match the throw speeds and throw angles required to reliably hit targets in the 30° Rotated Roll-to-Target tasks.

### Experiment Protocol

#### Experiment 1: Effects of visual cues and perturbation types on context-based flexibility of motor learning

Participants in Experiment 1 were divided into one of 4 experiment groups: Cued Rotated (n = 20, 11 female, 20.10 ± 2.17 years of age), Uncued Rotated (n = 21, 11 female, 20.05 ± 2.56 years of age), Cued Accelerated (n = 20, 14 female, 20.85 ± 5.89 years of age), or Uncued Accelerated (n = 19, 13 female, 21.05 ± 6.65 years of age); or one of 4 follow-up groups: Cued Rotated-15° (n = 15, 10 female, 19.80 ± 1.08 years of age), Uncued Rotated-15° (n = 14, 7 female, 21.00 ± 4.61 years of age), Cued Curved (n = 20, 12 female, 21.05 ± 4.31 years of age), or Uncued Curved (n = 20, 11 female, 20.35 ± 4.60 years of age) (Fig 1d).

After completing practice trials, an experimenter reiterated the task objectives. These instructions were also displayed within the virtual reality environment and participants were encouraged to mentally follow along (instructions found at osf.io/a5nv3/). The experiments were divided into multiple phases (Fig 1g). All experiments began with a Baseline phase where participants performed 40 Aligned Roll-to-Target tasks. These tasks were performed in trial-sets of 4 trials within which participants rolled the ball to each of the 4 possible targets and experienced each of the 4 possible Baseline Accelerations. The order of the target locations and Baseline Accelerations were randomized independently of each other within the trial-sets. The visual task surface was always aligned with the floor plane during the Baseline phase. Participants in the Cued and Uncued Aligned and Rotated groups performed an additional 40-trial Baseline phase using their non-dominant hands. This data was not analyzed for this study but is included in the data repository (osf.io/a5nv3/).

Following the Baseline phase, participants were asked to take a short break for a minimum of 1 minute. The experimenter then read further instructions informing the participants that although the environment may change, their goal was to always hit the target with the ball as accurately as possible. Participants in all groups then completed the Training phase (80 trials) where they performed one of 4 variations of the Roll-to-Target tasks, determined by their assigned group. The visual task surface was slanted by 25° (CCW roll) in all Cued groups and was aligned with the real-world floor plane in all Uncued groups. During the Training phase, the colour of the task surface was changed from black to green in the Cued groups in the four initial experiment groups, while the colour of the surface was changed to green for all groups in the follow-up groups.

To test the decay of learned behaviour when returning to baseline-like conditions, participants completed a Washout phase consisting of 40 trials of the Roll-to-Target task immediately following the Training phase. For all groups, the visual task surface was aligned with the floor plane during the Washout phase.

Participants in the Cued and Uncued Aligned and Rotated groups performed an additional 40-trial Transfer phase with their non-dominant hands. The Transfer phase was identical to the Training phase. This data was not analyzed for this study but is included in the data repository (osf.io/a5nv3/).

#### Experiment 2: Effects of visual surface-slant cues on implicit motor learning

In a second experiment (1 group: n=20, 14 female, 22.40 ± 5.88 years of age), we tested whether internal model-updating during motor adaptation relies on visual environmental cues – namely the visual slant of the surface in the Roll-To-Target tasks.

In this experiment we introduced Pivots: animations of the virtual task-surface played between phases of the experiment (Fig 1e, right side). Each Pivot lasted 2 seconds and involved a counter-clockwise (CCW) rotation of task surface on the yaw axis, either by 180° or 360°. During the Pivot animations, participants were verbally instructed via a recording played in the head mounted display system to disengage any cognitive strategies they may have used during the Training phase. Pivots were always followed by 8 Aligned Roll-to-Target trials to test the effect of the Pivot on performance (Fig 1h).

Participants first completed a Baseline phase, performing 2 sets Aligned Roll-to-Target tasks (16 trials, then 8 trials) separated by a 180° Pivot. Participants then repeated the 2 sets of Aligned Roll-to-Target task, separated by a 360° Pivot.

During the Training phase, participants performed 80 Rotated Roll-to-Target trials with a rotation magnitude of 30°, and the visual task surface was slanted by 25° (CCW roll), as in the Cued groups in Experiment 1. The colour of the task surface was changed to green during the Training phase.

During Test phases, we used 8-trial blocks to test the effect of a 180° or a 360° pivot of the task surface. We repeated the test blocks so that participants performed 4 test blocks following a 180° pivot, and 4 test blocks following a 360° pivot. Participants performed top-up Training blocks (20 trials of the Cued Rotated Roll-to-Target task) in between each set of test blocks (Fig 1h).

### Data Analysis

For each trial, we calculated the angle of the throwing motion in the Action step relative to a straight-line to the target along the task surface plane (termed “Throw Angle”), the speed of the throw, and the minimum distance to the target along the ball’s travel path. The Throw Angle was our primary measure of interest as participants were required to modify it to successfully hit targets in all our perturbed conditions. During Rotated and Curved Roll-to-Target tasks, changes in Throw Angles alone determined task success, while the speed of the throw also impacted task success in Accelerated Roll-to-Target tasks. We also calculated Minimum Errors for each trial: the smallest distance from any point on the ball path during the Feedback step to the centre of the target.

To account for individual differences in baseline throwing characteristics, we calculated the median Throw Angles of each participant when throwing to each of the 4 possible target locations during the Baseline Phase. We subtracted these target-specific baseline Throw Angles from the calculated Throw Angles in the Training and Washout phases. We similarly determined baseline errors for each target by calculating the median magnitude of error experienced during the Feedback step of the Roll-to-Target task. To obtain a measure of additional error due to our imposed perturbations, we subtracted these target-specific errors from the errors experienced during the Training and Washout phase.

To compare across groups, we normalized each participant’s Throw Angles relative to their median Throw Angles during the middle of the Training phase (trials 33-40). We then fit exponential decay functions to the change in each participant’s normalized Throw Angles during the first 40 trials of the Training and Washout phases.

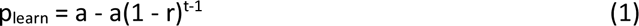

Equation 1 represents the change in performance over trials (t) in the Training phase (p_learn_) given a participant’s learning (r) and the asymptote of adaptation (a).

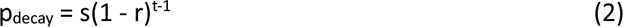

Equation 2 represents the decay in learned behaviour over trials (t) in the Washout phase (p_decay_) given a participant’s start point of decay (s) and decay rate (r). We assumed all participants started at baseline-like performance at the start of the Training Phase and returned to baseline-like performance at the end of the Washout phase. To visualize the reduction in errors over time during the Training phase, we also fit the exponential decay formula (Eq 2) to the changes in Minimum Errors over time in the Training phase.

We analysed the fit parameters for the start points and decay rates during the Washout phase (Eq 2) to determine if adaptation to our imposed perturbations led to the updating of existing models of motor control or the creation of new models. In instances of model-creation, we expected performance to return to baseline conditions quickly once a context change was detected during the Washout phase, signalled by low initial start points or fast decay rates. Alternatively, in instances of model updating, we expected high start points and relatively slower decay of learned behaviour during the Washout phase.

In Experiment 1, we tested whether a visual environmental cue or the perturbation type affected if internal models of motor control were updated. We used a 2x2 between-subject design where each participant experienced one of the 4 possible permutations of Visual Cue (Cued or Uncued) and Perturbation type (Accelerated or Rotated) combinations during the Training phase.

In Experiment 1, we first determined if the presence of the visual cue or perturbation type affected the characteristics of participants’ learning curves. We compared learning rates and asymptotes in the Training phase by conducting two 2x2 between-subject Analysis of Variance (ANOVA) tests with the presence of the visual cue and perturbation type as between-subject factors. When effects were found we also calculated and reported an effect size (generalized η^2^) and computed and reported inclusion bayes factors between models that included and did not include relevant effects [35]. We used Tukey’s HSD for post hoc pairwise analyses when necessary.

To determine if participants could flexibly change motor behaviour when experiencing previously seen contexts, we then compared start points and decay rates during the Washout phase with two additional 2x2 between-subject ANOVAs with the presence of the visual cue and perturbation type as between-subject factors. We again calculated generalized η^2^, inclusion bayes factors and Tukey’s HSDs when effects were found.

We repeated these 4 2x2 between-subject ANOVAs for Follow-up Experiment 1 and 2. In Follow-up Experiment 1, we compared Cued and Uncued Rotated-15 groups with the original Cued and Uncued Accelerated groups. In Follow-up Experiment 2, we compared the original Cued and Uncued Rotated groups with Cued and Uncued Curved groups.

In Follow-up experiment 1, we conducted additional 2x2 ANOVAs, with visual cue and perturbation type as between subject factors, comparing start points and error reduction rates fit to the reduction in Minimum Error during the Training Phase. We conducted two sets of 2x2 ANOVAs, first comparing the start points and error reduction rates of Minimum Error in the Accelerated groups to those in the Rotated groups, and second comparing the start points and decay rates of Minimum Error in the Accelerated groups to those in the Rotated-15 groups.

In Experiment 2, we tested whether environmental changes affected model updating. To elicit the updating of existing internal models, the Training phase was identical to the Cued Rotated group. We fit exponential decay functions (Eq 2) to the decay of learned behaviour during the 8-trial test phases. To determine if environment changes led to implicit performance changes, we then compared changes in Throw Angles during Test blocks following 180° pivots against those after 360° pivots using two paired t-tests. For all tests, we reported Bayes factors and Cohen’s d when effects were found.

## Results

### Experiment 1: Effects of visual cues and perturbation types on context-based flexibility of motor learning

We first compared Learning Rates and Asymptotes of Learning during the Training phase for all groups. When adapting to the imposed perturbations during the Training phase, participants in all groups altered their Throw Angles in a clockwise direction (Fig 2a-b). The presence of a visual environmental cue led to faster learning (mean_learning rate(cued)_ = 0.486, mean_learning rate(uncued)_ = 0.210; main effect of cue: F(1, 76) = 27.06, p < 0.001, η^2^ = 0.26, BFincl = 7.99x10^3^; Tukey HSD < 0.001, d = 1.16; Fig 2c-e). Adapting to Accelerated perturbations led to slightly higher asymptotes of adaptation (main effect of perturbation: F_(1, 76)_ = 4.09, p = 0.047, η^2^ = 0.051, BFincl = 0.996; mean_asymptote(accelerated)_ = 1.10, mean_asymptote(rotated)_ = 1.03. This effect however was weak and did not survive multiplicity correction (Tukey HSD =0.050, d = 0.452; Fig 2e: right plot). Although weak, we controlled for this effect in a follow-up experiment using Curved perturbations (see Follow-up experiment 2). Overall, the slant of the task surface acted as an informative visual cue when adapting to both Accelerated and Rotated perturbations, allowing for more rapid motor adaptation.

**Figure 2:**
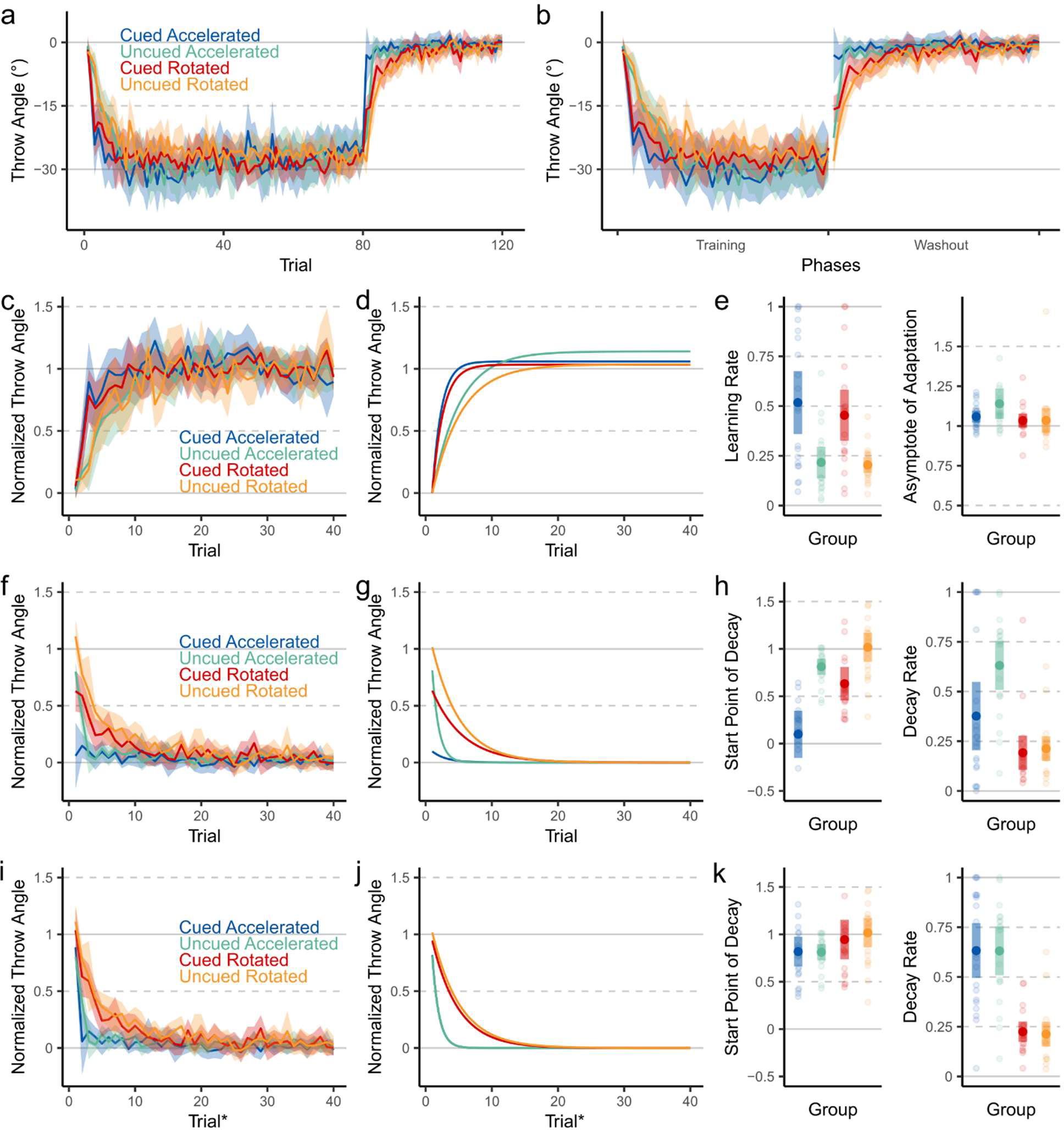
**a**: Throw angle deviations from straight-to-target throws (“Throw Angle”), relative to individual baseline Throw Angles, over the Training and Washout phases in Experiment 1. **b**: Throw Angles during the first 40 trials of Training and Washout phases. These trials were used for fitting exponential decay models. **c**: Normalized Throw Angles during the first 40 trials of the Training phase. **d**: Exponential decay functions fit to the changes in Normalized Throw Angles over trials in the Training phase. **e**: Learning rates and asymptotes of adaptation of fit Normalized Throw Angles of participants. **f**: Normalized Throw Angles during the Washout phase. **g**: Exponential decay functions fit to the changes in Normalized Throw Angles over trials in the Washout phase. **h**: Start points of decay and decay rates of Normalized Throw Angles of participants in the Washout phase. **i**: Normalized Throw Angles during the Washout phase including trials prior to cue detection (Trials+) in all Cued groups. **j**: Exponential decay functions fit to the changes in Normalized Throw Angles over Trials+ in the Washout phase **k**: Start points of decay and decay rates of Normalized Throw Angles of participants over Trials+ in the Washout phase. Shaded areas represent 95% confidence intervals.

To test the effects of the visual cues and perturbation types on model updating behaviour, we tested the speed at which participants could return to baseline-like performance levels during a Washout Phase, where all feedback was as in the Baseline Phase. We fit exponential decay functions to the decay of learned behaviour, i.e., the change in Normalized Throw Angles, when returning to unperturbed conditions. The Start Points were affected by both the presence of the visual cue (main effect of cue: F(1, 76) = 29.87, p < 0.001, η^2^ = 0.282, BF_incl_ = 2.16x10^4^) and the perturbation type (main effect of perturbation: F(1, 76) = 9.83, p = 0.002, η^2^ = 0.115, BF_incl_ = 21.69; Fig 2h: left plot). Uncued perturbations (mean_start point(cued)_ = 0.328, mean_start point(uncued)_ = 0.936; Tukey HSD < 0.001, d = -1.22) and Rotated perturbations (mean_start point(accelerated)_ = 0.458, mean_start point(rotated)_ = 0.807; Tukey HSD = 0.002, d = -0.702) both led to higher retained deviations in Throw Angles at the start of the Washout phase. The low amounts of retained Throw Angle deviation in the Cued Accelerated group was notably different from all other groups (Tukey HSD < 0.001 when compared pairwise to all other groups) and likely drove a significant interaction between the effects of perturbation type and the presence of the visual cue (F(1, 76) = 4.35, p = 0.040, η^2^ = 0.054, BFincl = 6.34).

The decay rates of learned behaviour during the Washout phase were also affected by both the presence of the visual cue (main effect of cue: F(1, 76) = 6.03, p = 0.016, η^2^ = 0.074, BF_incl_ = 4.18) and the perturbation type (main effect of perturbation: F(1, 76) = 29.01, p < 0.001, η_2_ = 0.276, BF_incl_ = 1.29*10^4^) (Fig 2h: right plot). Cued perturbations (mean_decay rate(cued)_ = 0.284, mean_decay rate(uncued)_ = 0.422; Tukey HSD = 0.016, d = -0.55) and Rotated perturbations (mean_decay rate(accelerated)_ = 0.504, mean_decay rate(rotated)_ = 0.203; Tukey HSD < 0.001, d = 1.21) led to lower decay rates of the learned change in Throw Angles during the Washout phase. Additionally, we found an interaction (F(1, 76) = 4.39, p = 0.040, η^2^ = 0.055, BFincl = 4.84) that was likely driven by the fast decay rates found in the Uncued Accelerated groups (Tukey HSD < 0.05 when compared pairwise to all other groups).

The surface slant cue affected both the Start Points and Decay Rates during the washout of learned behaviour. To determine if this cue served to initiate a context switch, we reanalyzed washout dynamics when including plausible context-switch detection in all groups. That is, we assumed that for the “Cued” groups, the change in surface slant during the first trial of the Washout phase would be sufficient to cue a context change, whereas, for the “Uncued” groups, the visual errors experienced during the first trial of the Washout condition would be necessary. To better compare the two groups, we included performance during the last trial of the Training phase for the “Cued” groups, ensuring our modelled Washout behaviour included context changes in all groups. We define Trials+ to be the relabelled trial numbers used for plotting and model-fitting. For all subsequent analyses, we used Trials+ to analyse the decay of motor adaptation during the Washout phase. Analyses of the uncorrected decay behaviour can be found in this project’s Open Science Framework repository (osf.io/a5nv3/).

When corrected for the detection of context changes, only the perturbation type affected the Start Points of decay functions fit to washout behaviour (main effect: F_(1, 76)_ = 4.76, p = 0.032, η^2^ = 0.059, BF_incl_ = 1.34; Fig 2i-k). Accelerated perturbations led to lower Start Points of decay (mean_start points (accelerated)_ = 0.815) than Rotated perturbations (mean_start points (rotated)_ = 0.981, Tukey HSD = 0.0323, d = -0.488; Fig 2k: left plot). Differences in the solution space between the perturbation types may have led to the differences in the Start points of decay. We explore this possibility in Follow-up Experiment 2. Perturbation type alone determined the decay rates of learned deviations of Throw Angle (main effect of perturbation type: F_(1, 76)_ = 75.59, p < .001, η^2^ = 0.499, BF_incl_ = 1.49*10^10^; Fig 2k: right plot). Adaptation to accelerated perturbations decayed more rapidly than adaptation to rotated perturbations (mean_decay rate (accelerated)_ = 0.632, mean_decay rate (rotated)_ = 0.219; Tukey HSD < 0.001, d = 1.95.). The decay of adaptation to accelerated perturbations was fast, taking on average 2 trials to return to baseline-like performance, suggesting motor learning that is flexible when context changes are detected. Following adaptation to rotated perturbations, the lower decay rates suggest existing internal models were updated.

### Follow up experiment 1: Rotated 15° perturbations

For each trial, we defined Minimum Errors as the shortest distance between any point on the ball’s travel-path during the Feedback phase and centre of the target. Although we designed our task to match the adaptation rates of Throw Angles during the Training phase for both Accelerated and Rotated groups, participants in the Accelerated groups experienced smaller Minimum Errors during the Training phase (Fig 3a-b). When fitting Eq 2 to Minimum Errors experienced in the Training phase, participants in the Accelerated groups experienced lower initial Minimum Errors (main effect of perturbation on starting points: F(1, 76) = 57.00, p < 0.001, η^2^ = 0.429, BF_incl_ = 9.52*10^7^; mean_initial minimum error(accelerated)_ = 1.67, mean_initial minimum error(rotated)_ = 3.37; Tukey HSD < 0.001, d = -1.69) and reduced Minimum Error more quickly (main effect of perturbation on learning rate: F_(1, 76)_ = 7.05, p = 0.010, η^2^ = 0.085, BFincl = 3.75; mean_decay rate(accelerated)_ = 0.244, mean_decay rate(rotated)_ = 0.125; Tukey HSD = 0.010, d = 0.594; Fig 3b-c). To determine if differences in Minimum Error reduction led to the choice between model updating or flexible motor learning, we conducted a follow-up experiment matching the Minimum Error experienced between the two perturbation types. To do so, we repeated the Cued and Uncued Rotated conditions with a second pair of Rotated conditions using a 15° rotation to the Throw Angle: the “Rotated-15” groups (Fig 1d-e: dark red and dark yellow).

**Figure 3:**
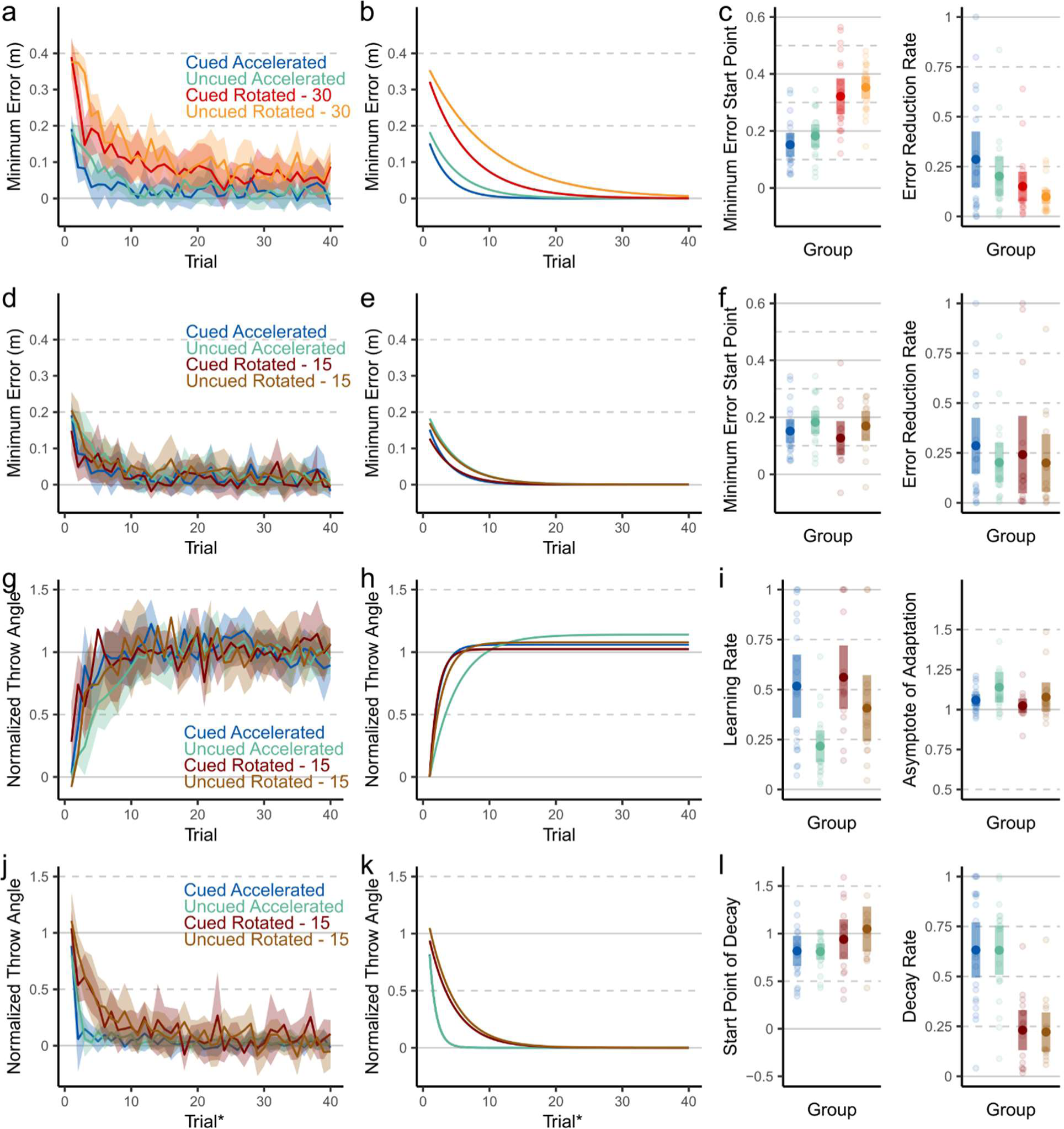
Minimum Errors during Training phases and Throw Angles during Training and Washout phases. **a**: Error experienced during the Training phase in Accelerated and Rotated groups. **b**: Exponential decay functions fit to the reduction of error over trials in the Training phase in Accelerated and Rotated groups. **c**: Start points and decay rates of fit Minimum Errors experienced by participants in Accelerated and Rotated groups. **d**: Error experienced during the Training phase in Accelerated and Rotated-15 groups. **e**: Exponential decay functions fit to the reduction of experienced error over trials in the Training phase in Accelerated and Rotated-15 groups. **f**: Start points and decay rates of fit Minimum Errors experienced by participants in Accelerated and Rotated-15 groups. **g**: Normalized Throw Angles during the Training phase in Accelerated and Rotated-15 groups. **h**: Exponential decay functions fit to the changes in Normalized Throw Angles over trials in the Training phase in Accelerated and Rotated-15 groups. **i**: Learning rates and asymptotes of adaptation of fit Normalized Throw Angles of participants in Accelerated and Rotated-15 groups. **j**: Normalized Throw Angles during Trials+ in the Washout phase in Accelerated and Rotated-15 groups. **k**: Exponential decay functions fit to the changes in Normalized Throw Angles over Trials+ in the Washout phase in Accelerated and Rotated-15 groups. **l**: Start points of decay and decay rates of Normalized Throw Angles of participants during Trials+ in Accelerated and Rotated-15 groups. Shaded areas represent 95% confidence intervals.

To ensure both the Accelerated groups and the Rotated-15 groups experienced a similar evolution of Minimum Errors over time, we again fit Eq 2 to Minimum Error over the first 20 trials of the Training phase (Fig 3d-e). When adapting to 15° rotations, initial Minimum Errors were more likely under models that did not include effects of perturbation type, cue, or interactions between the two (BF_incl_ = 0.258, 0.539, 0.147, respectively; Fig 3f: left plot). Similarly, the rates of Minimum Error reduction between the Accelerated groups and the Rotated-15 groups were more likely under models that did not include effects of perturbation type, cue, or interactions between the two (BF_incl_ = 0.190, 0.265, 0.076, respectively; Fig 3f: right plot). After confirming that error reduction rates and starting errors were similar across all groups, we repeated the analyses in Experiment 1 using normalized Throw Angles.

During the Training phase, learning rates were affected by the presence of the visual cues (main effect of cue: F(1, 64) = 11.30, p < 0.001, η^2^ = 0.150, BFincl = 32.84; Fig 3h-i). Cued perturbations led to a faster learning rate (mean_learning rate(cued)_ = 0.540, mean_learning rate(uncued)_= 0.312; Tukey HSD = 0.001, d = 0.824; Fig 3i: left plot). This was driven by participants in the Uncued Accelerated group learning slower than both the Cued Accelerated and Cued Rotated-15 groups (Fig 3i: left plot). We found large individual differences in the learning rates of the Uncued Rotated-15 group, which was not reliably different from any other groups during post-hoc comparisons. The Asymptotes of Adaptation were similar across all groups; the observed asymptotes were more likely under models that did not include an effect of perturbation type, or an interaction effect between perturbation type and the presence of a surface slant cue (BF_incl_ = 0.474, 0.303, respectively; the effects of the presence of the surface slant cue were inconclusive (BF_incl_ = 1.140); Fig 3i: right plot).

We used Trials+ to analyze the decay behaviour of learned Throw Angle deviations during the Washout phase of this follow-up experiment (Fig 3j-k), accounting for when participants detected a context change. The Start Points of the decay functions were weakly affected by the perturbation type (main effect: F_(1, 64)_ = 4.98, p = .029, η^2^ = 0.072, BF_incl_ = 1.44; Fig 3i: left plot). Accelerated perturbations led to lower Start Points compared to Rotated perturbations (mean_start points (accelerated)_= 0.815, mean_start points (rotated-15)_ = 0.994; Tukey HSD = 0.029, d = -0.547). In line with our initial results, only the perturbation type affected the rates of decay (main effect: F_(1, 64)_ = 47.66, p < .001, η^2^ = 0.427, BF_incl_ = 3.74*10^6^), with Accelerated perturbations leading to faster decay of adaptation (mean_decay rate (accelerated)_ = 0.632, mean_decay rate (rotated-15)_ = 0.227; Tukey HSD < 0.001, d = 1.69: Fig 3i: right plot). Our primary findings from Experiment 1 remained consistent when adjusting for the sizes of experienced errors during the Training phase. This suggests that the differences in the propensity for model updating was not influenced by the observed differences in error sizes, but the types of the perturbation experienced.

### Follow-up experiment 2: Curved perturbations

When adapting to Accelerated perturbations, participants could hit the target with the thrown ball through specific combinations of Throw Angles and Throw Speeds. Figure 4a shows the solution space, the combinations of Throw Angle and Throw Speed used during the Training phase of the experiments and their associated Minimum Error sizes, for each target when adapting to Accelerated perturbations. The solution manifold is the sub-space within the solution space that led to direct target hits, indicated by white-coloured data points. The solution manifolds for Accelerated perturbations were a function of both Throw Angle and Throw Speed (Fig 4a), whereas the solution manifolds for Rotated perturbations was a function of only the Throw Angle, with some added noise due to the Baseline Accelerations (Fig 4b). Additionally, a larger range of Throw Angles could lead to task success when adapting to Accelerated perturbations (Fig 4a). To determine if these solution space differences during the Training phase affected our findings, we conducted a second follow-up experiment where the ball-path during the Feedback step of the Roll-to-Target task was curved (see Fig 1f). The ball path was deviated from a straight-to-target path by an amount proportional to the distance from the starting position, so that when at the target distance, the angular deviation relative to the starting position is always 30°. Curved perturbations resulted in solution manifolds like Rotated perturbations (Fig 4b-c) while retaining visual feedback properties of Accelerated perturbations (Fig 1f). Once we ensured similar solution manifolds for both perturbation types, we compared changes in Throw Angles in the Curved groups with those in the Rotated groups to determine if our findings from Experiment 1 were driven by the different solution manifolds.

**Figure 4:**
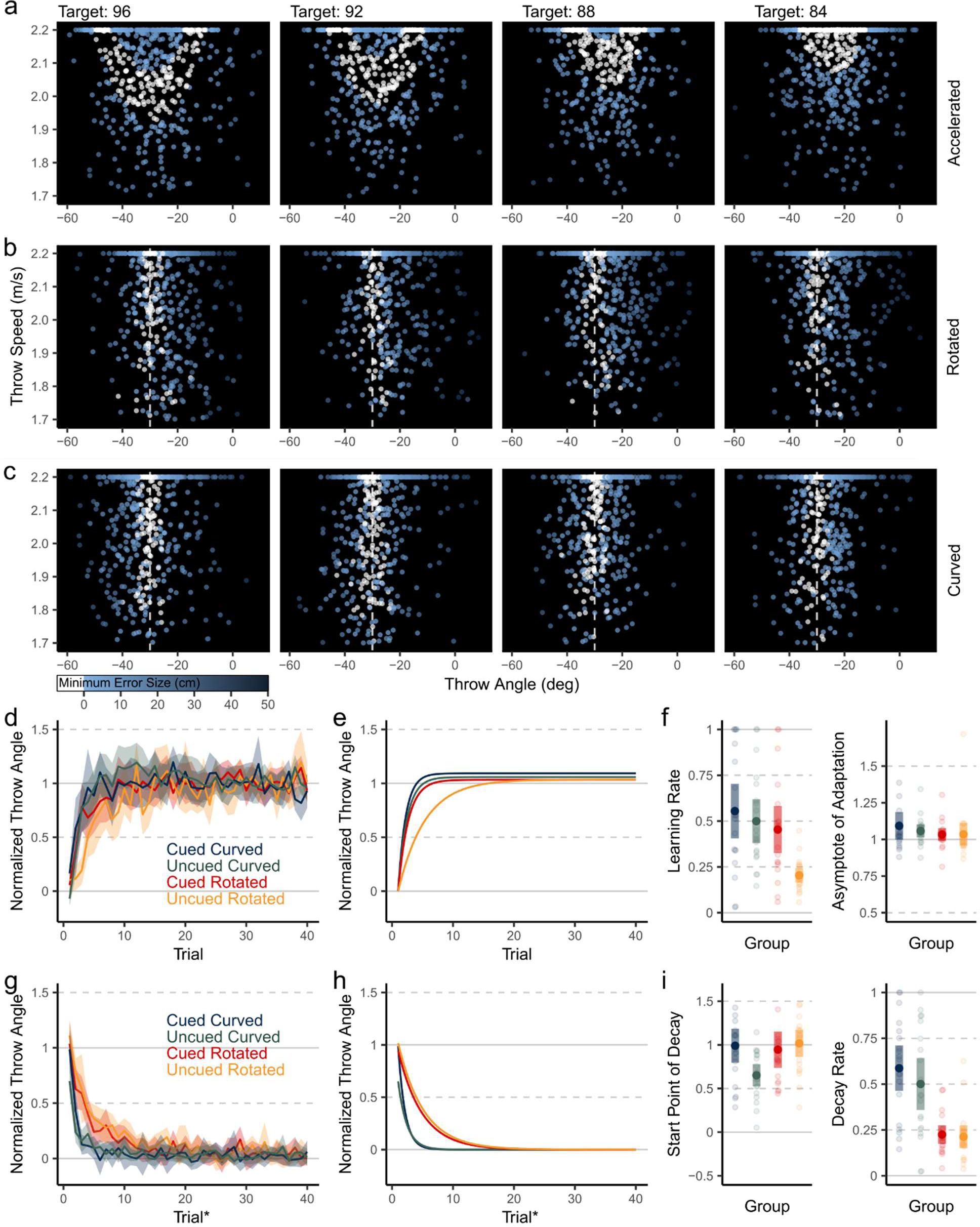
Minimum error sizes experienced (where white data points indicate a direct target hit) during throws in the Feedback step during the Training phase in Accelerated (**a**), Rotated (**b**), and Curved (**c**) groups as a function of Throw Angles and throw speeds in the Action step. **d**: Normalized Throw Angles during the Training phase in Curved and Rotated groups. **e**: Exponential decay functions fit to the changes in Normalized Throw Angles over trials in the Training phase in Curved and Rotated groups. **f**: Learning rates and asymptotes of adaptation of fit Normalized Throw Angles of participants in Curved and Rotated groups. **g**: Normalized Throw Angles during Trials+ in the Washout phase in Curved and Rotated groups. **h**: Exponential decay functions fit to the changes in Normalized Throw Angles over Trials+ in the Washout phase in Curved and Rotated groups. **i**: Start points of decay and decay rates of Normalized Throw Angles of participants during Trials+ in Curved and Rotated groups. Shaded areas represent 95% confidence intervals.

During the Training phase, learning rates were affected by both the perturbation type (main effect: F_(1, 77)_ = 12.88, p < .001, η^2^ = 0.143, BF_incl_ = 55.74) and the presence of the visual cue (main effect: F_(1, 77)_ = 7.60, p = .007, η^2^ = 0.090, BF_incl_ = 7.61; Fig 4e, f: left plot). Participants in the Uncued Rotated groups learned notably slower than all other groups (Tukey HSD < 0.010, pairwise against all groups), likely driving the observed effects (Fig 3f: left plot, orange data). The asymptotes of adaptation were similar across all groups (BF_incl_ = 0.329, 0.191, 0.090 for perturbation type, presence of visual cue, and an interaction, respectively; Fig 4f: right plot).

We again analysed the decay of learned behaviour with respect to the detection of context changes using Trials+. Participants in the Uncued Curved group had lower Throw Angles prior to the induced decay of adaptation during the Washout phase when compared to those in the Cued Curved group (Tukey HSD = 0.027) and the Uncued Rotated group (Tukey HSD = 0.013), driving an interaction effect between the perturbation type and the presence of a visual cue (interaction: F_(1, 77)_ = 6.10, p = .016, η^2^ = 0.073, BF_incl_ = 2.82; Fig 4i: left plot). Like our findings in Experiment 1 and Follow-up Experiment 1, only the perturbation type affected the decay rates (main effect of perturbation type: F_(1, 77)_ = 44.94, p < .001, η^2^ = 0.369, BF_incl_ = 2.63*10^6^; Fig 4i: right plot). The performance changes of adaptation to Curved perturbations decayed at a faster rate than those to Rotated perturbations (mean_decay rate (curved)_ = 0.544, mean_decay rate (rotated)_ = 0.219; Tukey HSD < 0.001, d = 1.49.). The visual perturbation type, that is, acceleration-like perturbations, and not the differences in solution spaces during the Training phase in Experiment 1, determined the propensity for internal model updating.

### Experiment 2: Effects of visual surface slant cues on implicit motor learning

Once we understood that Rotated perturbations likely led to model updating, we tested whether information about the environment informed the internal models being updated. In a second experiment, participants adapted again to a 30° rotation perturbation to Throw Angles, cued by a visual slant of the task surface as in the Rotated-Cued group in Experiment 1. Whereas the Washout phase in the Cued-Rotated group was cued by a visually flat surface on which to roll the ball, in this experiment, multiple Test phases were cued by an animation that pivoted the task surface either 180°, leading to a surface slant opposite that in the Training phase, or 360°, leading to the same surface slant as in the Training phase (Fig 1h). As expected, since internal models must be reupdated during the Test phases, the pivot angle did not have an effect on decay rates during the Test phases (t_(19)_ = 09.59, p = 0.564, BF_10_ = 0.27; Fig 5b: right plot). However, 180° animations led to less negative start points of decay (mean_180°_ = -10.90, mean_360°_ = -19.70; t_(19)_ = -4.16, p < 0.001, d = 1.23, BF_10_ = 63.44) than 360° animations (Fig 5b: left plot). Crucially, participants were asked to not use cognitive strategies during the Washout phases in Experiment 2. These findings suggest that the internal models updated when adapting to Rotated perturbations include environment characteristics like the slant of the surface that can lead to flexible motor behaviour.

**Figure 5:**
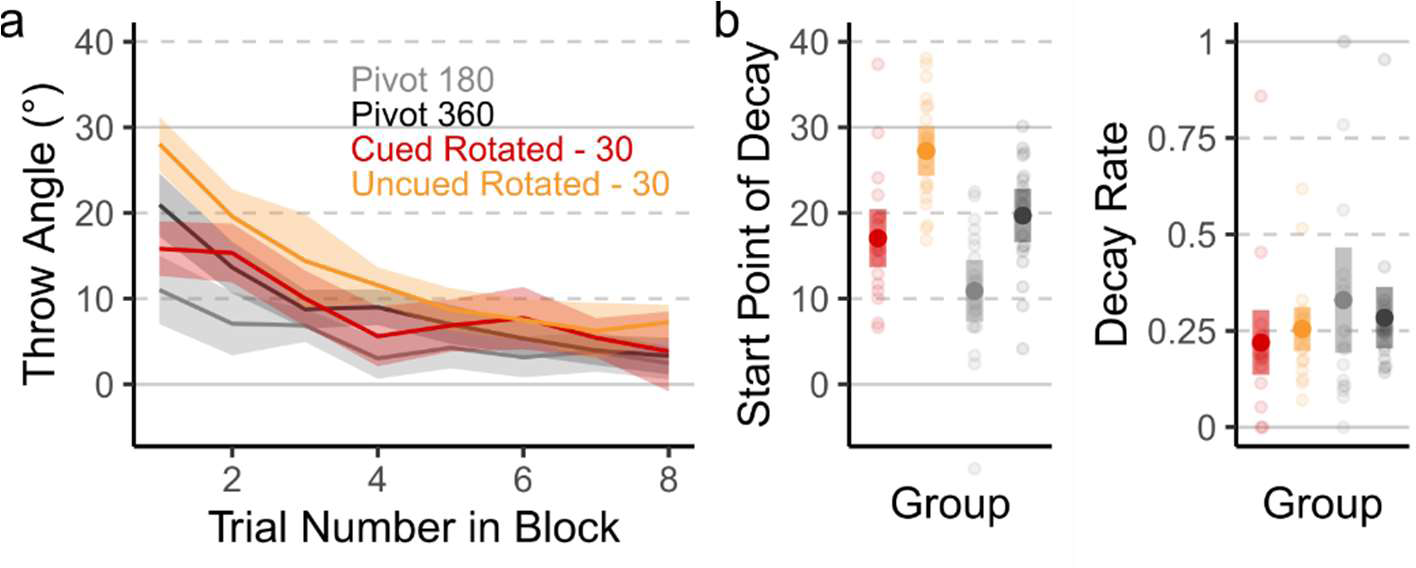
**a**: Changes in Throw Angles over 8-trial Test phases in the Pivot Experiment, along with the first 8 trials in the Washout phase of the Cued and Uncued Rotated 30° groups from Experiment 1. **b**. Start points of decay and decay rates during 8-trial Pivot 180 and Pivot 360 Washout phases, along with start points of decay and decay rates during the first 8 trials of the Washout phase of the Cued and Uncued Rotated 30° groups from Experiment 1. Shaded areas represent 95% confidence intervals.

## Discussion

In our virtual ball-throwing task, the tendency for model updating during motor adaptation was entirely determined by the type of perturbation experienced. When errors could be plausibly attributed to object-environment interactions, as in real-world-like accelerations of ball paths, motor learning was flexible, and performance could quickly switch between recently learned and previously experienced contexts. When the visual feedback of the hand-ball interactions at the moment of release were misaligned instead, suggesting internal sources of error, we observed evidence for the updating of internal models of motor control. Immersive visual cues about the task environment that reliably predicted motor-context switches enabled faster initial learning and fast switching to existing internal models but did not facilitate the creation of new motor memories. Although changes in the immersive virtual environment during perturbations did not determine if models were updating during motor learning, properties of the virtual environment, like the surface slant, did inform the implicit internal models involved in executing a target-directed ball-rolling action.

In our study, Accelerated perturbations led to motor adaptation that could be flexibly changed when previously experienced contexts were detected. Accelerated perturbations can plausibly be explained by physical processes acting on the rolling ball, such as the effects of gravity when rolling on a slanted surface or the effects of sidewinds when rolling on a flat surface. Participants may have intuitively simulated physics properties that govern interactions in the environment [36,37] to explain away unpredicted behavior and supress model updating [26]. Alternatively, recent findings suggest that people fail to accurately estimate accelerations while still compensating for acceleration-based errors over time [38]. It would then follow that acceleration-like errors, like the Curved perturbations in our follow up experiment that could not be explained by Newtonian mechanics, should also lead to learning similar to acceleration based perturbations. That is indeed what we observed. Surprisingly, visual explanations for the acceleration perturbation, that is, the slant of the surface, did not have any additional effects on the tendency for model updating, suggesting visual information did not provide any additional information for the assignment of error to an external source.

Although motor changes during adaptation were matched across Accelerated and Rotated groups, participants in the Rotated groups experienced larger errors during trials in the early Training phase. Larger errors may also lead to slower decay, both due to higher amounts of implicit adaptation, and models being re-updated taking longer to return to baseline. Learning rates should be similar across learning rates when errors are similarly reliable [8–10], but large error sizes could lead to large explicit changes in performance [39]. Findings in our follow-up study using 15° visuomotor rotations suggest the perturbation type, and not experienced error sizes, determined if internal models were updated. Although experienced errors in the 30° Rotated group were large, they may not have been large enough to elicit the use of large strategies that could be suppressed or changed in response to a changes in the external environment [39,40].

A successful real-world, target-directed throw can be achieved by various combinations of throwing speeds and throwing directions due to redundancies in the task. The solution subspace for our Accelerated tasks, i.e., a manifold along the solution space that led to successful task outcomes, allowed for higher variability in throwing directions than the Rotated conditions. When task solutions allow for redundancy, variability is only reduced to a limited degree [41,42]. A wide solution subspace may allow for both increased motor exploration along the throwing angle dimension as well as reduced reinforcement learning to throwing directions specific to the perturbation, allowing for faster return to baseline-like performance [43–45]. However, since adaptation to Curved perturbations where the solution space was identical to the Rotated perturbations was flexible, our findings suggest that visually observed properties of the perturbation, and not the solution space determine if models are updated.

In our follow-up experiments, although the colour of the task surface changed during the training phase, only the visual change in the task-surface slant reliably cued an immediate change in motor performance. Our findings support previous work suggesting colour changes in the environment are ineffective at eliciting dual learning, where the a creation of multiple context specific motor memories is necessary [18,20,27]. Generally, properties of the task environment that do not directly affect movement dynamics, termed “indirect cues”, do not reliably lead to the creation of new context-specific motor memories [18,20,23]. In our study however, both the visual slant and colour of the task surface were indirect cues containing no direct information about dynamic body states during a throwing movement. Our findings from Experiment 1 and Experiment 2 suggest that environment properties serving as indirect cues that explain the behaviour of interactable objects, as opposed to less task-relevant cues, are effective at prompting the development of both explicit and implicit changes in behaviour.

Although prior works suggest likely strong contributions of explicit learning processes to visual slant-related learning [27], in this study, we do not distinguish whether a lack of model updating was due to the creation of cognitive strategies to counter the perturbation, or the creation of new implicit motor memories. Indeed, neither the use of explicit strategies during motor learning nor implicit motor learning processes prevent the parallel development of either class of adaptation [39,40,46,47]. In Experiment 2, we directly tested for implicit components of surface-slant specific adaptation. When participants were asked to exclude any strategies while throwing to targets on a slanted surface, we observed implicit slant-specific motor learning. Since we relied on verbal instructions, there is a chance that participants did not disengage explicit slant-specific aiming strategies. However, in such cases, we expect throws to the opposite direction when the task surface was pivoted 180°. Instead, all but one participant reduced their trained throwing angles in response to 180° pivots while maintaining the learned deviation direction, suggesting that the observed slant-specific behaviour changes were indeed implicit.

Overall, our results suggest that adaptation to acceleration-like perturbations to the path of a thrown object is flexible and does not rely on updating internal models of motor control. Additionally, informative task relevant environment cues may facilitate immediate and implicit behaviour changes but do not affect the propensity for internal model updating during motor learning.

## Acknowledgements

This work was supported by the Natural Sciences and Engineering Research Council of Canada (NSERC), and Vision: Science to Application (VISTA) for S.M; and NSERC for D.Y.P.H. The funders had no role in study design, data collection and analysis, decision to publish, or preparation of the manuscript.

## Author Contributions

S.M., B.M.tH., and D.Y.P.H designed the experiments. S.M., and A.K. conducted the experiments. S.M., analyzed the data. S.M. wrote the manuscript and prepared figures. All authors reviewed and approved the manuscript.

## Ethics Declarations

The authors declare no competing interests.

## References

1. Miall RC, Wolpert DM. Forward models for physiological motor control. Neural Networks. 1996;9. doi:10.1016/S0893-6080(96)00035-4

2. Shadmehr R, Krakauer JW. A computational neuroanatomy for motor control. Experimental Brain Research. 2008. doi:10.1007/s00221-008-1280-5

3. Shadmehr R, Smith MA, Krakauer JW. Error Correction, Sensory Prediction, and Adaptation in Motor Control. Annu Rev Neurosci. 2010;33: 89–108. doi:10.1146/annurev-neuro-060909-153135

4. Krakauer JW, Ghilardi MF, Ghez C. Independent learning of internal models for kinematic and dynamic control of reaching. Nat Neurosci. 1999. doi:10.1038/14826

5. Leow LA, Gunn R, Marinovic W, Carroll TJ. Estimating the implicit component of visuomotor rotation learning by constraining movement preparation time. J Neurophysiol. 2017;118: 666– 676. doi:10.1152/jn.00834.2016

6. Haith AM, Huberdeau DM, Krakauer JW. The influence of movement preparation time on the expression of visuomotor learning and savings. J Neurosci. 2015. doi:10.1523/JNEUROSCI.3869-14.2015

7. Smith MA, Ghazizadeh A, Shadmehr R. Interacting adaptive processes with different timescales underlie short-term motor learning. PLoS Biol. 2006;4: 1035–1043. doi:10.1063/1.2184639

8. Baddeley RJ, Ingram HA, Miall RC. System identification applied to a visuomotor task: Near-optimal human performance in a noisy changing task. J Neurosci. 2003;23. doi:10.1523/jneurosci.23-07-03066.2003

9. Wei K, Körding K. Uncertainty of feedback and state estimation determines the speed of motor adaptation. Front Comput Neurosci. 2010;4. doi:10.3389/fncom.2010.00011

10. van Beers RJ. How Does Our Motor System Determine Its Learning Rate? PLoS One. 2012;7. doi:10.1371/JOURNAL.PONE.0049373

11. Kunavar T, Cheng X, Franklin DW, Burdet E, Babič J. Explicit learning based on reward prediction error facilitates agile motor adaptations. PLoS One. 2023;18: e0295274. doi:10.1371/JOURNAL.PONE.0295274

12. McGarity-Shipley MR, Heald JB, Ingram JN, Gallivan JP, Wolpert DM, Flanagan JR. Motor memories in manipulation tasks are linked to contact goals between objects. J Neurophysiol. 2020;124: 994–1004. doi:10.1152/jn.00252.2020

13. Ingram JN, Howard IS, Flanagan JR, Wolpert DM. Multiple Grasp-Specific Representations of Tool Dynamics Mediate Skillful Manipulation. Curr Biol. 2010;20: 618–623. doi:10.1016/j.cub.2010.01.054

14. Ahmed AA, Wolpert DM, Flanagan JR. Flexible Representations of Dynamics Are Used in Object Manipulation. Curr Biol. 2008;18. doi:10.1016/j.cub.2008.04.061

15. Kong G, Zhou Z, Wang Q, Kording K, Wei K. Credit assignment between body and object probed by an object transportation task. Sci Rep. 2017;7. doi:10.1038/s41598-017-13889-w

16. Ayala MN, ‘t Hart BM, Henriques DYP. Concurrent adaptation to opposing visuomotor rotations by varying hand and body postures. Exp Brain Res. 2015;233: 3433–3445. doi:10.1007/s00221-015-4411-9

17. Ghahramani Z, Wolpert DM. Modular decomposition in visuomotor learning. Nature. 1997;386. doi:10.1038/386392a0

18. Howard IS, Wolpert DM, Franklin DW. The effect of contextual cues on the encoding of motor memories. J Neurophysiol. 2013;109: 2632–44. doi:10.1152/jn.00773.2012

19. Seidler RD, Bloomberg JJ, Stelmach GE. Context-dependent arm pointing adaptation. Behav Brain Res. 2001;119. doi:10.1016/S0166-4328(00)00347-8

20. Baldeo R, Henriques DYP. Dual adaptation to opposing visuomotor rotations with similar hand movement trajectories. Exp Brain Res. 2013;227: 231–241.

21. Woolley DG, Tresilian JR, Carson RG, Riek S. Dual adaptation to two opposing visuomotor rotations when each is associated with different regions of workspace. Exp Brain Res. 2007;179: 155–165.

22. Hinder MR, Woolley DG, Tresilian JR, Riek S, Carson RG. The efficacy of colour cues in facilitating adaptation to opposing visuomotor rotations. Exp Brain Res. 2008;191: 143–155. doi:10.1007/s00221-008-1513-7

23. Gandolfo F, Mussa-Ivaldi FA, Bizzi E. Motor learning by field approximation. Proc Natl Acad Sci USA. 1996;93: 3843–3846. doi:10.1073/pnas.93.9.3843

24. Osu R, Hirai S, Yoshioka T, Kawato M. Random presentation enables subjects to adapt to two opposing forces on the hand. Nat Neurosci. 2004;7: 111–112.

25. Hegele M, Heuer H. Implicit and explicit components of dual adaptation to visuomotor rotations. Conscious Cogn. 2010;19: 906–917. doi:10.1016/j.concog.2010.05.005

26. Scarfe P, Glennerste A. Humans use predictive kinematic models to calibrate visual cues to three-dimensional surface slant. J Neurosci. 2014;34: 10394–10401. doi:10.1523/JNEUROSCI.1000-14.2014

27. Forano M, Schween R, Taylor JA, Hegele M, Franklin DW. Direct and indirect cues can enable dual adaptation, but through different learning processes. J Neurophysiol. 2021;126: 1490–1506. doi:10.1152/jn.00166.2021

28. Hinder MR, Tresilian JR, Riek S, Carson RG. The contribution of visual feedback to visuomotor adaptation: How much and when? Brain Res. 2008;1197: 123–134. doi:10.1016/j.brainres.2007.12.067

29. Stirk JA, Underwood G. Low-level visual saliency does not predict change detection in natural scenes. J Vis. 2007;7: 1–10. doi:10.1167/7.10.3

30. Spotorno S, Faure S. Change detection in complex scenes: Hemispheric contribution and the role of perceptual and semantic factors. Perception. 2011;40. doi:10.1068/p6524

31. Heald JB, Ingram JN, Flanagan JR, Wolpert DM. Multiple motor memories are learned to control different points on a tool. Nat Hum Behav. 2018;2: 300–311. doi:10.1038/s41562-018-0324-5

32. Sternad D, Huber ME, Kuznetsov N. Acquisition of novel and complex motor skills: Stable solutions where intrinsic noise matters less. Adv Exp Med Biol. 2014. doi:10.1007/978-1-4939-1338-1_8

33. Levac DE, Huber ME, Sternad D. Learning and transfer of complex motor skills in virtual reality: A perspective review. J Neuroeng Rehabil. 2019;16: 1–15. doi:10.1186/s12984-019-0587-8

34. Brookes J, Warburton M, Alghadier M, Mon-Williams M, Mushtaq F. Studying human behavior with virtual reality: The Unity Experiment Framework. Behav Res Methods. 2020;52: 455–463. doi:10.3758/s13428-019-01242-0

35. Clyde MA, Ghosh J, Littman ML. Bayesian adaptive sampling for variable selection and model averaging. J Comput Graph Stat. 2011;20: 80–101. doi:10.1198/jcgs.2010.09049

36. Battaglia PW, Hamrick JB, Tenenbaum JB. Simulation as an engine of physical scene understanding. Proc Natl Acad Sci U S A. 2013;110: 18327–18332. doi:10.1073/pnas.1306572110

37. Smith KA, Vul E. Sources of Uncertainty in Intuitive Physics. Top Cogn Sci. 2013;5: 185–199. doi:10.1111/tops.12009

38. Brenner E, Rodriguez AI, Muñoz VE, Schootemeijer S, Mahieu Y, Veerkamp K, et al. How Can People Be so Good at Intercepting Accelerating Objects if They Are so Poor at Visually Judging Acceleration? Iperception. 2016;7: 1–13. doi:10.1177/2041669515624317

39. Bond KM, Taylor JA. Flexible explicit but rigid implicit learning in a visuomotor adaptation task. J Neurophysiol. 2015. doi:10.1152/jn.00009.2015

40. Modchalingam S, Vachon CM, ’t Hart BM, Henriques DYPP, Hart BM t., Henriques DYPP. The effects of awareness of the perturbation during motor adaptation on hand localization. PLoS One. 2019;14. 10.1371/journal.pone.0220884

41. Huber ME, Kuznetsov N, Sternad D. Persistence of reduced neuromotor noise in long-term motor skill learning. J Neurophysiol. 2016;116. doi:10.1152/jn.00263.2016

42. Cohen RG, Sternad D. Variability in motor learning: Relocating, channeling and reducing noise. Exp Brain Res. 2009;193. doi:10.1007/s00221-008-1596-1

43. Uehara S, Mawase F, Therrien AS, Cherry-Allen KM, Celnik P. Interactions between motor exploration and reinforcement learning. J Neurophysiol. 2019;122. doi:10.1152/jn.00390.2018

44. Haith AM, Krakauer JW. Model-Based and Model-Free Mechanisms of Human Motor Learning. Adv Exp Med Biol. 2013;78. doi:10.1016/j.earlhumdev.2006.05.022

45. Zhang Z, Guo D, Huber ME, Park S-W, Sternad D. Exploiting the geometry of the solution space to reduce sensitivity to neuromotor noise. 2018. doi:10.1371/journal.pcbi.1006013

46. Albert ST, Jang J, Modchalingam S, ’t Hart M, Henriques D, Lerner G, et al. Competition between parallel sensorimotor learning systems. Elife. 2022;11. 10.7554/eLife.65361

47. Miyamoto YR, Wang S, Smith MA. Implicit adaptation compensates for erratic explicit strategy in human motor learning. Nat Neurosci. 2020;23: 443–455. doi:10.1038/s41593-020-0600-3

